# Impact of Population on Yayo Biosphere Reserve, Ilubabor Zone, Oromia Regional State, Southwest Ethiopia

**DOI:** 10.1101/2025.04.26.650807

**Authors:** Fikru Mosisa, Adanech Asfaw, Tefera Jogora

## Abstract

The research was carried out in the Yayo Coffee Forest Biosphere Reserve, located in the Oromia region of southwest Ethiopia. Data were gathered from key informants, kebele leaders, development agents, selected households, focus group discussions, and field observations. Land use and land cover change data were obtained from satellite imagery spanning 1984 to 2007. Population growth data were sourced from the Ethiopian Central Statistical Agency’s Jimma branch. Additionally, field visits included direct observations of the forest’s condition. A total of 69 participants responded to both structured and semi-structured questionnaires designed to address the study’s objectives. The communities in the study area do not rely on a single livelihood strategy but instead engage in various activities such as crop cultivation, forestry, livestock rearing, and off-farm work, with varying levels of dependence on each. Among the forest products collected, coffee, firewood, honey, and wild spices were the most common. The main factors contributing to forest degradation were ranked as population growth (88.6%), illiteracy (75.7%), lack of local participation (73.9%), poverty (64.3%), investment (47.1%), and urbanization (42.9%), in decreasing order. Satellite image analysis revealed a negative correlation between population growth and forest cover, with a significant difference at P<0.05. The study highlights the impact of population pressure on forest resource degradation in the Yayo Coffee Forest Biosphere Reserve, stressing the need for effective strategies focused on sustainable management, conservation, and utilization of the biosphere reserve.

## INTRODUCTION

Forests are a valuable resource that offers numerous benefits to human well-being. The effects of demographic changes on forests and the environment are frequently examined through the concept of biological carrying capacity, which refers to the maximum number of individuals that a resource can support. However, various factors can impact carrying capacity, including economic development, socio-political processes, trade, technology, and consumption patterns[1].

The primary causes of land degradation are deforestation, overgrazing, excessive logging, shifting cultivation, and poor agricultural practices concerning soil and water management. These include the failure to implement soil and water conservation techniques, improper crop rotation, use of marginal lands, incorrect application of fertilizers, mismanagement of irrigation systems, and over-extraction of groundwater. Indirect causes of land degradation include population growth, land scarcity, insecure or short-term land tenure, as well as poverty and economic pressures.[2, 3]. In Ethiopia, alongside rapid population growth and low socio-economic development, land degradation has significantly impacted the country’s ecological balance. In the early 20th century, Ethiopia’s forest cover accounted for 40% of its total land area; however, today, this has dwindled to less than 2.2%.[4]. The larger portions of the existing forests are even secondary forest [1]. Despite receiving some attention, deforestation remains a significant issue in the Buffer and Transitional zones, requiring urgent solutions. This study aims to examine the impact of population growth on the natural forests within the Yayo Biosphere Reserve.

## Materials and methodology

### Study area description

The Yayo Coffee Forest Biosphere Reserve is located in the Ilu Abba Bora Zone of the Oromia Regional State in southwestern Ethiopia. It is recognized as the center of origin for Coffea arabica, the world’s most widely consumed coffee. Yayo is the largest and most significant forest for conserving wild coffee populations globally. The reserve plays a crucial role in preserving both natural and cultural landscapes. Situated in the southwestern part of the Oromia region, the Yayo Biosphere Reserve includes the Woredas of Hurumu, Yayo, Chora, Nopha, Alge Sachi, and Doreni, covering coordinates from 8°0’42” to 8°44’23” N and 35°20’31” to 36°18’20” E.

The district’s elevation ranges from 1,139.2 meters to 2,581.9 meters above sea level, with the lowest point located at Gaba River and the highest point at the summit of Sayi Mountain (2,581 meters) in Keresi. The area experiences a hot and humid climate, with an average annual temperature of around 23°C, ranging from a mean minimum of 18.59°C to a mean maximum of 27.88°C (YDRADO, 2018). The varied physical conditions and altitudinal differences contribute to a rich diversity of climate, soil, and vegetation, fostering the development of numerous plant species with a high level of diversity[5]. The rainfall pattern of the districts varies annually from 1,191.6 to 1,960.7mm showing variations from year to year. It is a unimodal type of rainfall that increases from May to October and declines in November.

This study focuses on three districts Hurumu, Yayo, and Doreni within the Yayo Coffee Forest Biosphere Reserve, which represent three distinct climatic zones. The area is composed of 3.5% highland (5,750.4 hectares), 85% temperate zone (138,465.85 hectares), and 11.47% lowland (18,684.75 hectares) [6, 7]. The diverse climatic conditions and habitats in these districts have contributed to a high level of species diversity in both plants and animals. This biodiversity richness is one of the reasons why Ethiopia is considered one of the 20 most biodiverse countries in the world (Fig1).

**Figure 1:**
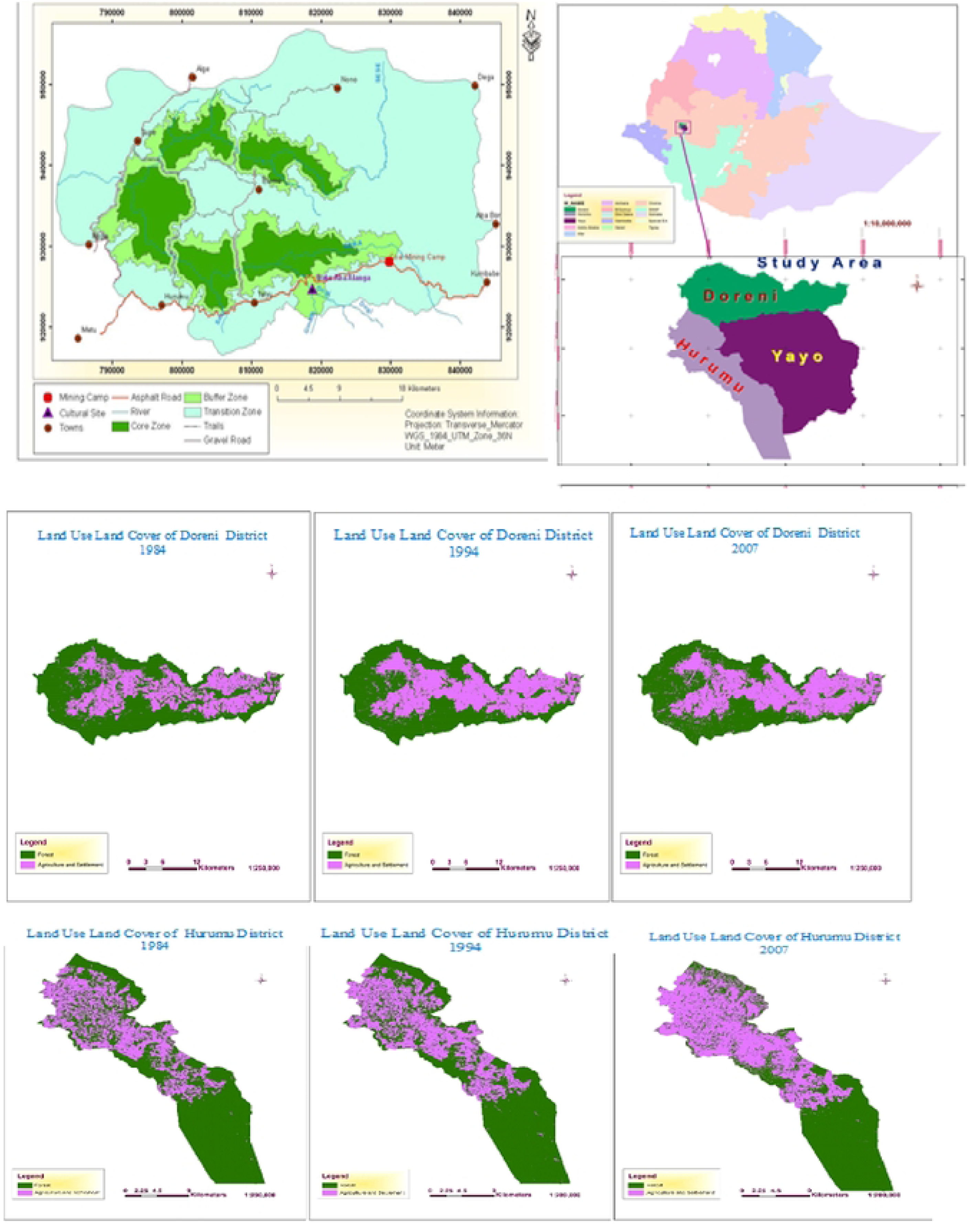
Maps of the study area (Sources: Ethiopian ArcMap data using GIS software)

### Data collection

The data for this study includes a sample household survey, household attributes, socio-economic characteristics, and land use/land cover change data. The research approach involves a consultative and interactive process, engaging respondents who are willing to provide essential information. Interviews were conducted to collect data on the role of local communities in forest conservation, the impacts of human activities on the forest, perceptions toward conservation efforts, as well as the challenges and consequences associated with these activities.

To collect data, formal survey questionnaires were used to gather quantitative information from selected households. Focus group discussions and interviews were conducted to collect primarily qualitative data. Satellite imagery was utilized to generate land use/land cover change data. Additionally, field visits were carried out, complemented by direct observations of the forest conditions.

### Study Population

Interviews were conducted with local community members, local leaders, and representatives of community social and economic groups. Additionally, key informants, including wildlife and forest experts from the Ilu Ababora Zonal and Woreda levels, were also interviewed. Furthermore, two focus group discussions were held with local community members in the sampled areas to gather more insights.

### Sample Size and Sampling Techniques

A preliminary survey was conducted to gain an overview of the Yayo Coffee Forest Biosphere Reserve’s distribution and to select sample woredas and nearby kebeles for the study. Out of the six woredas in the biosphere (Yayo, Hurumu, Chora, Bilo Nopha, Alge Sachi, and Doreni), three were chosen as sample woredas: Yayo, Hurumu, and Doreni. From these three woredas, nine kebeles Haro, Gaba, Wangegne, Waboo, Geci, Wixete, Boco, Badesa, and Henna were randomly selected. Three kebeles were chosen from each woreda, with a total of nine adjacent rural kebeles selected purposefully. Respondents were selected based on their proximity to the Yayo Coffee Forest, the length of time they have lived in the forest reserve area, and their local knowledge of forest resource conservation.

A total of 69 general respondents were participated in the household survey data collection. In addition, 18 key informants (2 from each kebele), including major stakeholders such as village leaders, kebele leaders, development agents (DAs), and experts, were interviewed. Furthermore, three focus group discussions were conducted with seven participants in Hurumu, seven in Dorani, and eight in Yayo woreda, respectively. Generally 109 respondents were participated during data collection processes.

### Data Analysis

The collected data were reviewed, corrected, and coded using Microsoft Excel. Qualitative data were interpreted thematically, while quantitative data were analyzed using descriptive statistics and econometric methods. The analysis was conducted using the Statistical Package for Social Sciences (SPSS) version 20.

## Results and Discussion

### Socio-economic characteristics

During the study time, there were a total of 91,694 households in the region found unevenly distributed over the then 3 woredas and with a total population of 458,472 individuals (Table 1). The majority of the sample households (97.1%) were male-headed, while only 2.9% were female-headed. The ages of the respondents ranged from 20 to 80 years, with a mean age of 48 years. Among the sampled households, 92.8% were married, and 7.2% were single, with no respondents being widowed or divorced. Of the male household heads who were married, about 95.7% had one wife, while the remaining 4.3% had more than one wife. The family size varied between 1 and 12 members, with an average of 5.18, which is consistent with the national average family size. Regarding education, 15.9% of the respondents were illiterate (unable to read or write), 30.4% had attended grades 1-5, 42% had completed grades 6-12, and 11.6% had obtained a diploma or higher education, up to the first degree (Table 1). Similarly, the family Socio-economic Status includes the household income, earners’ education and occupation as well as combined income when their own attributes are assessed [8, 9]. In fact, the socio-economic status can be measured in a number of different ways. Most commonly, it is measured by father’s education, occupation and income [10].

**Table 1:** Back ground of the households by sex, age, education level and marital status Back ground.

| Back ground | Distinction | Frequency | Percent (%) |
| --- | --- | --- | --- |
| Sex | Male | 67 | 97.1 |
|  | Female | 2 | 2.9 |
|  | Total | 69 | 100.0 |
| Age | 20-30 | 2 | 2.9 |
|  | 31-40 | 27 | 39.1 |
|  | 41-50 | 16 | 23.2 |
|  | 51-60 | 10 | 14.5 |
|  | >61 | 14 | 20.3 |
|  | Total | 69 | 100.0 |
| Educational status | Illiterate | 11 | 15.9 |
|  | Grade 1-5 | 21 | 30.4 |
|  | Grade 6-12 | 29 | 42.0 |
|  | Diploma & above | 8 | 11.6 |
|  | Total | 69 | 100.0 |
| Marital status | Married | 64 | 92.8 |
|  | Single | 5 | 7.2 |
|  | Total | 69 | 100.0 |

### Livelihood strategies

In this study, among the sample households that participated in the survey, 8.7% reported that their sole income source was forest product collection, 2.9% relied solely on crop production, and 1.4% depended exclusively on off-farm activities. The result is in line with [11]. The larger the household size of the respondent, the higher was the likelihood of access to forest. Meanwhile, 37.7% of respondents indicated that their income came from a combination of crop production, forest products, and livestock production. Against above result, 73% forest dwellers depend on these products as a source of revenue in the WAJIB, Bale Zone, southern Ethiopia [12]. Additionally, 27.5% reported that their income sources were crop production and forest products, 5.8% relied on crop and livestock production, 4.4% depended on forest products and off-farm activities, and 2.9% reported that their income was derived from a mix of crop production, forest products, and off-farm activities. The remaining 8.7% of respondents indicated that they relied on all types of household income-generating activities (Table 2). The value of income sources like forest products, crop production, livestock production and off-farm activates depend on each other [13]. Research conducted in the adjacent district of yayo, included under one NFPA with *Gabba*-*Dogi* i.e., Yayo NFPA [14], revealed that 92.6 percent of the population in the study area have coffee in the forest [14] from which 57.3 kg of honey on average is harvested per household per year (Ibid:).

**Table 2:** The main livelihood strategies in the Yayo Coffee Forest Biosphere Reserve.

| No. | Livelihood activities | Frequency | Percentage |
| --- | --- | --- | --- |
| 1 | Crop production only | 2 | 2.9 |
| 2 | Forest product collection only | 6 | 8.7 |
| 3 | Off-farm activities only | 1 | 1.4 |
| 4 | Crop and livestock | 4 | 5.8 |
| 5 | Crop and Forest product | 19 | 27.5 |
| 6 | Crop, livestock and forest product | 26 | 37.7 |
| 7 | Crop, forest product and off-farm | 2 | 2.9 |
| 8 | Forest products and off-farm activities | 3 | 4.4 |
| 9 | All livelihood activities | 6 | 8.7 |
| Total |  | 69 | 100 |

### Institution acting as stakeholders in the Yayo Coffee Forest Biosphere

#### Formal Institutions in forest management

Based on the information gathered from interviews and focus group discussions, various institutions at the federal, regional, and local levels were identified, along with their interlinkages concerning the management of the coffee forest resource under investigation.

Changes in the institutional structure of the Ministry of Agriculture (MoA) since the early 1990s have failed to establish a dedicated government body for the management, conservation, and sustainable use of coffee forests. This gap has contributed, to some extent, to the increasing lack of effective institutions for the sustainable management of coffee forests. Federal institutions, under the [8] mainly institution for Biodiversity Conservation (IBC), provide technical support for Yayo coffee forest Biosphere Reserve conservation.

At the regional level, two institutions with potential connections to the coffee forest are examined: the Oromia Forest and Wildlife Enterprise Supervising Agency and the regional ARDB. The primary issues identified within these organizations include a lack of technical and direct focus on forest coffee biodiversity conservation, insufficient budget and technical personnel at the ARDB, inadequate decentralization of the budget, and limited community involvement in planning and execution at the Forest Enterprise Supervising Agency. Another gap noted in the State Forest Enterprise is the absence of incentives to motivate local communities to conserve specific forests. The study by [15]confirms the gradual marginalization of local indigenous institutions with the increasing control of state that enforced formal institutions under different regimes during the past.

At the local level, the institutions involved in the use, conservation, and management of the coffee forest include the Yayo Coffee Forest Biosphere Reserve Conservation Project, the district administration (comprising the kebele and development team), and the district ARDO.

#### Informal institutions contributing for forest management

The study examines various informal institutions in terms of their structure and role in the livelihoods of the community in the study area. These institutions are categorized into four groups, with particular emphasis placed on two clusters. The first cluster consists of territorial-based administrative indigenous/customary institutions, while the second includes a variety of self-help work organizations. The territorial-based administrative indigenous/customary institutions are further divided into four groups: Tuullaa, Xuxee, Shane, and Jaarsa Biyya, along with Muchoo. The study conducted by [16, 17] revealed the same result with current study.

Tuullaa, Xuxee, and Shane can contribute to coffee forest management in two key ways. First, Tuullaa, which organizes, leads, and enforces the activities, rules, and regulations of various local customary institutions and self-help work organizations, plays a critical role. These customary institutions are directly or indirectly involved in coffee forest management, including overseeing the harvesting of coffee from the forest and enforcing rules to prevent violations in the management of the coffee forest [18]. Ostrom in this regard stated that stronger norms demonstrated by reciprocity may be needed to protect the commons [19].

Secondly, since Tuullaa is central to the overall social, cultural, and economic life of the community in the coffee forest area, it can also regulate the activities and behavior of the local population. Specifically, it plays a key role in promoting collective action by facilitating the design, development, and enforcement of rules and alternative institutions that encourage sustainable coffee forest management through active community participation [20]

On the other hand, Jaarsa Biyya and Muchoo can contribute to the conservation and management of the coffee forest, as well as other natural resources, in two main ways. First, they can enforce the rules and regulations of existing customary institutions by serving as customary judges in natural resource management or conflict resolution cases. They have the authority to uphold any customary rules that the local community respects and adheres to. Second, these institutions have the potential to adjudicate matters within the local community and enforce additional rules related to coffee forest and other resource management, provided they are granted the authority to do so. With broad community acceptance, Jaarsa Biyya and Muchoo can actively participate in designing and enforcing rules that govern the sustainable management of the coffee forest ([16, 21]. Tesema also indicated that the existence of a variety of neighborhood voluntary self-help associations helped Oromo to generate surplus production, food security and self-sufficiency [22].

### Conservation policies and its drawbacks

As indicated by various informants in the coffee forest study area, different forms of ownership rights existed prior to the demarcation process. Portions of the coffee forests were owned by private individuals, the state (public ownership), associations, communities, or a combination of these entities. Oromia Rural Land Use and Administration Proclamation [23]; Forest Proclamation of Oromia, Proclamation ([24]) and Federal Forest Development, Conservation, and Utilization Proclamation no. [25, 26], have sure implication to natural resource management, even though in the practical implementation they have faced their own limitation.

### The contributions of the Biosphere reserve to the local communities

In this study, the collection of forest products was identified as the primary source of household income. Approximately 88.4% of respondents stated that they are highly dependent on forest products to support their households. Additionally, 52.2% of the households surveyed indicated that collecting forest products is their top priority for sustaining their livelihoods (see Table 3). This finding is similar with many studies conducted in different areas in Ethiopia. For instance [27]at Yayo district same study area,[28] at Sheka zone, [29], At Bale Mountain, forest products were revealed as the primary source of income, with contributions of 54%, 49%, and 44.7%, followed by crop production. In contrast, other studies have indicated that forest products contribute as the fourth most important source of livelihood for households. For instance, [30] at Liban wereda, Borena, south Ethiopia (32%), [31] at Gore district, southwest Ethiopia similar Agro-ecology (23%). While 29% of the respondents said forest products collections are serving my household as second income source and 13% of them have ranked forest products as third option for their livelihood (table3).

**Table 3:** Forest products priority to support the household’s livelihood.

| No. | Priority | No of respondent | Percent |
| --- | --- | --- | --- |
| 1 | Forest product is as 1<sup>st</sup> priority | 36 | 52.2 |
| 2 | Forest product is as 2<sup>nd</sup> priority | 20 | 29 |
| 3 | Forest product is as 3<sup>rd</sup> priority | 9 | 13 |
| 4 | Not engaged | 4 | 5.8 |
| 5 | Dependency level | 65 | 94.2 |
| Total |  | 69 | 100.0 |

### The major factors of forest depletion

The analysis of the household survey results highlighted several factors contributing to the depletion of forest stocks in the Yayo Coffee Forest Biosphere Reserve. Among the households surveyed, approximately 88.6% of respondents identified Urbanization as a major factor in the depletion of forest resources. This is due to the local population’s reliance on primary activities such as wood logging, non-timber forest product collection, and farming (see Table 4).

**Table 4:** Factors responsible for the possible depletion of forest stocks in the reserve.

| No. | Factors | Frequency | Percent |
| --- | --- | --- | --- |
| 1 | Urbanization | 62 | 88.6 |
| 2 | Timber extraction | 53 | 75.7 |
| 3 | Agricultural expansion | 45 | 64.3 |
| 4 | Investment | 33 | 47.1 |
| 5 | Climate change effects | 30 | 42.9 |
| 6 | Lack of local people participation | 51 | 73.9 |

Approximately 73.9% of respondents strongly agreed that the lack of indigenous people participation in the management and protection of the reserve is a major cause of forest depletion (see Table 4). Since local communities, who are the primary stakeholders, are not sufficiently involved in the reserve’s protection and management, a sense of ownership has not been fostered. Additionally, 75.7% of respondents identified Timber extraction as a contributing factor to the forest depletion. Around 64.3% of respondents also pointed to Agricultural expansion as a significant factor, driving local communities to rely on the forest for their immediate consumption needs (see Table 4).

On the other hand, 47.1% of respondents identified industrial investment as a major factor contributing to forest depletion. The Yayo Fertilizer Factory, for instance, has cleared large areas of natural forest for its operations. The factory has utilized 67 hectares of land at the Witate site for thermal energy plantations and 38 hectares at the Achebo site for coal mining, totaling 115 hectares of buffer zone land within the biosphere reserve. The majority of this land has been cleared of natural forest.

Additionally, the local community noted that the Yayo Fertilizer Factory has displaced many farmers from their land, forcing them into daily labor. According to the study, approximately 2,200 farming households were physically displaced, losing access to vital livelihood resources, including wild coffee farms, homes, and agricultural land, in order to make way for the factory’s operations [32] implementation. In fact the research team also witnessed more than twenty six (26) local poor household engaged in daily labor to support their livelihood. In this regard, the arrival of the COFCOP [32]. The coal mining program, launched in 2010, was initially estimated to create job opportunities for thousands of unemployed individuals from the area. According to an interview with the project manager of the Yayo Fertilizer Factory, between 2012 and 2013, the factory employed over 6,000 skilled and unskilled workers, many of whom were resettled in the highly forested villages of the district, particularly in Achebo and Wutete. However, most of these employees, ranging from low-level laborers to higher-ranking officials, were brought in from other regions, especially from Tigray, excluding the local community from the workforce.

Since 2018, however, the youth from the displaced households have been organized and have taken control of the coal mining activities at the Achebo site.

Regarding physical displacement, hundreds of village residents were denied access to their coffee farms, forest areas, and local water sources due to the strict security measures implemented by the project. While compensation was provided to those displaced, it was only for the loss of annual crops (such as maize, sorghum, and barley), perennial crops (like mango, pawpaw, and oranges), garden trees, and village houses and structures. However, the study revealed that compensation was calculated for only 420 of the 2,200 displaced individuals. For example, compensation for a coffee stand was estimated at just 6.50 birr, and for one hectare of farmland, it was only 400 birr.

The study found the compensation to be inadequate for several reasons. First, the compensation was calculated based on a fixed period, which failed to account for the long-term, intergenerational value of wild coffee. Second, the total compensation amount was far below the current market value and did not consider the cultural and social importance of the expropriated assets. Third, compensation payments were made directly to the affected individuals without providing any guidance or training on how to sustainably manage their livelihoods after displacement.

42.9% of respondents identified urbanization, particularly construction activities, as another factor contributing to the depletion of forest stocks (see Table 4). As a reminder, the Yayo natural forest canopy used to cover the main road to Yayo town, as shown in Figure 2, highlighting the extent of the forest loss due to development activities. Other study result indicated lack of economic opportunities and social amenities in rural areas, rural-urban migration has resulted in a considerable population increase in many sub-Saharan African cities [33].

**Figure 2:**
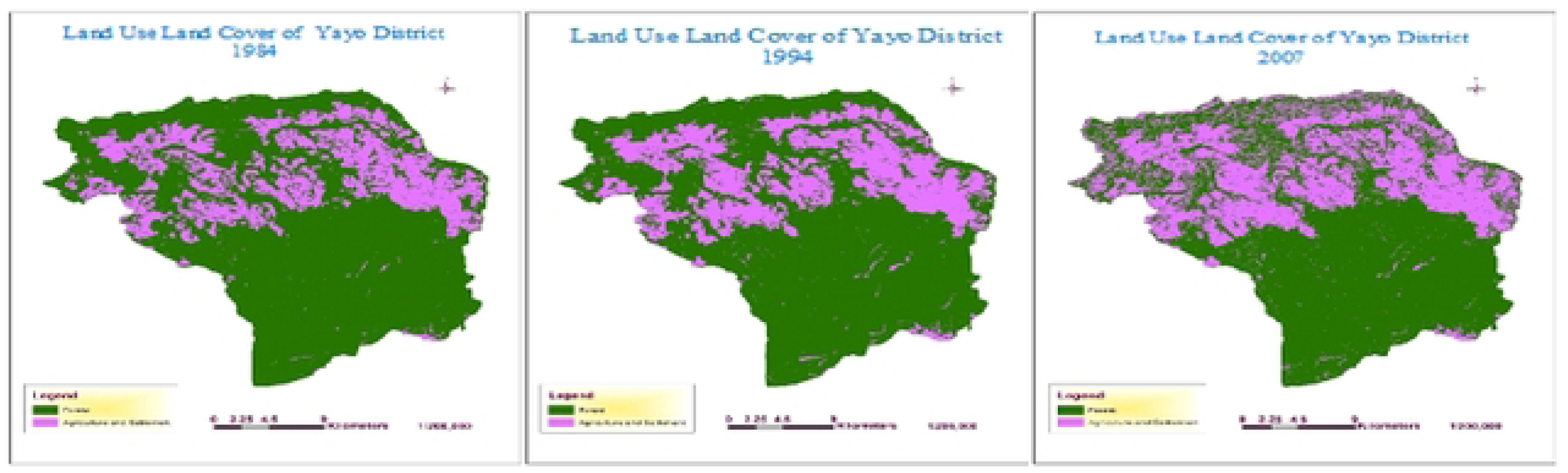
**Satellite Image of Land Use Land Cover (1984 to 2007)**

### Population growth and forest cover change

Population pressure in the study area results from two interconnected factors: natural population growth and migration, particularly the resettlement of people into forested areas. Recently, the area has seen a significant influx of migrant settlers, both in terms of numbers and ethnic diversity, arriving at various times and for different reasons. The majority of these migrants are from the Amhara ethnic group, particularly from Wello, Gojjam, and Gondar. Additionally, migrants from other regions and ethnic groups have also settled in the area, including people from the Southern Nations, Nationalities, and Peoples’ Region (SNNPR), such as the Kembatas, Hadiyas, and Guraghes; Tigrians from the north; and Oromos from the eastern part of Ethiopia.

As revealed by the statistical analysis from the household survey, approximately 88.6% of respondents identified population growth as a major factor contributing to the depletion of forest stocks in the biosphere reserve. This is because the local population engages with the forest for various livelihood needs. The primary reasons for this engagement include farmland expansion, overgrazing, settlement development, and the collection of forest products, all of which were identified as the dominant causes (see Table 5).

**Table 5:**
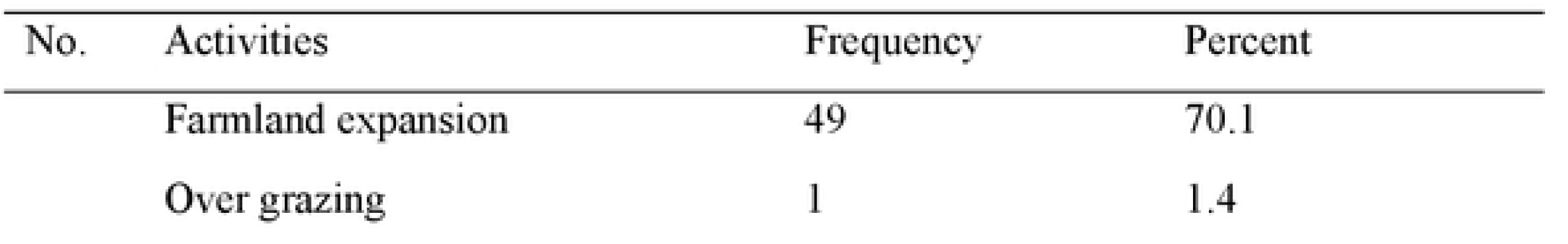
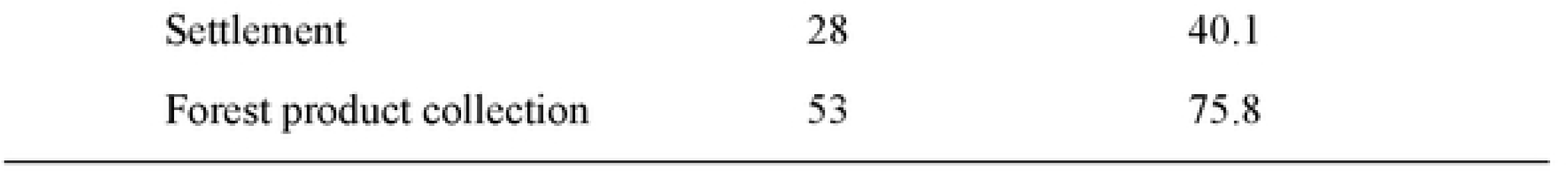
forest land conversion.

As shown in Table 5, the household survey data reveals that about 75.8% of participating households identified forest product collection as a major factor contributing to forest degradation, particularly forest coffee production and firewood collection. Key informants also provided additional insights, noting that in Hurumu Woreda, specifically in Gaba Kebele (two zones: Siso and Darimu) and Wangegne Kebele (two zones: Tije and Konchi), there is a concerning level of exploitation of trees, especially Cordia Africana and other species, for illegal timber production. This activity, which is highly destructive to the forest, was not captured in the household survey but remains a significant issue in the area.

Conversion of forest land into agricultural land reported as the second most important direct cause of deforestation by 70.1 % of the respondents and this result in line with study by different scholars [34–36]. According to information gathered from interviews and focus group discussions with local informants, farmers in the study area prefer planting khat (Catha edulis) by clearing forest areas for two main reasons. First, cultivating khat offers a comparative advantage over wild coffee, as it is more resilient to external challenges such as climate change and plant diseases. Second, khat leaves can be harvested throughout the year, allowing farmers to spread out their labor demands over the entire year. However, a perceived disadvantage of khat cultivation is that it involves higher production costs compared to wild coffee.

On the other hand, settlements were identified as the third major factor contributing to forest degradation, with 40.1% of respondents citing it as a cause (see Table 5). Analysis of responses from focus group discussions, key informants, and expert interviews suggests that the sharp increase in population in the district has created various social, economic, and political challenges at the local level.

Growing competition for land and resources has intensified pressures on forest biodiversity, leading to the degradation of Coffea arabica genetic material. Secondly, the influx of migrants from different cultural, technological, and institutional backgrounds has further complicated the situation. Many of these newcomers, unfamiliar with the local socio-ecological conditions, lacked the traditional ecological knowledge and management practices that had been developed by the indigenous communities over centuries. As these migrants focused on extracting economic benefits from the forests, they were not part of the established cultural forest management systems and were often unaware of the local ecology or concerned with the long-term protection of forest resources. As a result, they undermined the customary management practices that had been designed to ensure the sustainable use and preservation of forest biodiversity.

In this study area, overgrazing is not considered a serious problem for forest degradation, as the landholding systems in the area include designated grazing land (kaloo) specifically allocated for livestock, separate from forested areas (see Table 5).

### Population growth and forest cover change satellite image result

Population data for the past 40 years were obtained from the Central Statistical Agency (CSA) of Jimma Branch, based on the Ethiopian Population Censuses conducted in 1984, 1994, and projection of 2007 census. According to the census data, the population of Ilubabor Zone was over 847,048 in 1984, over 970,243 in 1994, and over 1,271,609 in 2007 projected to 2,301,242 in 2024, increased by 10.3%. The three districts—Yayo, Hurumu, and Dorani—selected as the study area within the six districts included in the biosphere reserve, had a combined population of 77,438 in 1984, 132,223 in 1994, and 248,811 in 2007 projected to 375,822 in 2024, which increased by 9.6% in average (see Table 6).

**Table 6:** Population growth data.

| Year | Population Growth in 1984, 1994 and 2007 |  |  |  | Total |
| --- | --- | --- | --- | --- | --- |
|  | Ilubabor Zone | Yayo Wereda | Hurumu | Durani |  |
| 1984 GC | 847,048 | 29,837 | 27,586 | 20,015 | 77438, |
| 1994GC | 970,243 | 52,851 | 42,667 | 36,705 | 132223 |
| 2007GC | 1,271,609 | 83,579 | 106,294 | 58,938 | 248811 |
| 2024 GC | 2,301,242 | 139,001 | 165,143 | 97,491 | 375822 |

The forest cover change in Yayo Biosphere Reserve over the past 40 years (from 1984 to 2024) is presented as Satellite images comparison Landsat 5 TM for 1984, Landsat 5 TM and ETM for 1994, and Landsat 7 TM and ETM+ for 2024, each with a 30-meter resolution satellite images cover showed Forest coverage decreased from 120087.2 hectares to 100772.9 hectares or by 11.6% over the 40-year period (see Figure 2). This finding is agreed with [37], Forest coverage is replaced by agricultural land, and settlement land are the dominant land use types in the study area, which is divided into three compartments: core, buffer, and transitional zones, within the Yayo Biosphere Reserve. LANDSAT/TM satellite images from 1986 to 1990 show that Ethiopia’s forest cover had since then been reduced to 3.93%, or 45,055 sqkm [38].

On the other hand, information gathered from household surveys, key informant interviews, and focus group discussions also confirmed that the expansion of agricultural land and settlements has significantly contributed to the decrease in forest cover within the Yayo Coffee Forest Biosphere Reserve. These land use changes were identified as key drivers of forest degradation in the region, further supporting the findings from satellite imagery and statistical analysis. Which is very dangerous for the biosphere reserve to conserve it’s Biodiversity that enabled to receive the Recognitions from [39]. According to information gathered during focus group discussions and key informant interviews with farmers in Yayo District, specifically in the Kori area near Sayi Forest, around 15 households have illegally settled on 8 hectares of the core zone of the biosphere reserve. Similarly, in Ilu Abba Dinka Kebele, Janeh area, about 28 households have settled on 14 hectares of the core zone. In total, approximately 22 hectares of forest in the core zone have been cleared due to agricultural expansion and coffee production. Additionally, in some areas, particularly in Gechi Kebele, local people have raised concerns about the planting of exotic tree species, which they believe could rapidly threaten and negatively affect the indigenous species by clearing the native vegetation. Similarly, studies by [40] in eastern Africa on three protected forest areas in the area demonstrated that deforestation and degradation of natural forests as a result of agricultural activities pose adverse impacts on the biodiversity of natural forest areas.

The present results align with a study conducted in the Hawa Galan district of Kelem Wollega, Ethiopia, which found that forest cover in the district was negatively correlated with both the district’s overall population and the population within the forest area. [41]. According to [42], also reported that population expansion has placed increasing stress on limited natural resources i.e. forests and woodland have been destroyed.

The Pearson Correlation analysis results indicate a strong negative correlation between forest cover and population growth rate over the past 40 years in the surveyed forest biosphere reserve, with a coefficient value of -0.998. This negative value confirms an inverse relationship between population growth and forest cover—meaning that as population growth increases, forest area decreases, and vice versa. The statistical analysis also shows that this correlation is significant at the 0.05 confidence level (P < 0.05) (see Table 8). In line with the above result, the works of [43] indicates the impacts and pressures of population on LULC changes are highly dependent on population density and Results of the Pearson Correlation for forest coverage (P = -0.006) were negatively associated with population pressure on forest.

**Table 7:** Satellite Images Results of Land Use Land Cover (1984 to 2007).

| LULC* | 1984 |  | 1994 |  | 2007 |  |
| --- | --- | --- | --- | --- | --- | --- |
|  | Area (Ha) | % | Area (Ha) | % | Area (Ha) | % |
| Doreni district |  |  |  |  |  |  |
| Forest | 29030.1 | 63.1 | 25395.59 | 55.2 | 24889.52 | 54.1 |
| Agriculture & Settlement | 16976.4 | 36.9 | 20610.91 | 44.8 | 21116.98 | 45.9 |
| Total | 46006.5 | 100 | 46006.5 | 100 | 46006.5 | 100 |
| Hurumu district |  |  |  |  |  |  |
| Forest | 30848.4 | 66.3 | 29638.65 | 63.7 | 24241.35 | 52.1 |
| Agriculture & Settlement | 15680.1 | 33.7 | 16889.85 | 36.3 | 22287.15 | 47.9 |
| Total | 46528.5 | 100 | 46528.5 | 100 | 46528.5 | 100 |
| Yayo district |  |  |  |  |  |  |
| Forest | 60208.67 | 74.5 | 57703.34 | 71.4 | 51642.06 | 63.9 |
| Agriculture & Settlement | 20608.34 | 25.5 | 23113.66 | 28.6 | 29174.94 | 36.1 |
| Total | 80817 | 100 | 80817 | 100 | 80817 | 100 |
| Forest total | 120087.17 | 69.3 | 112737.58 | 65 | 100772.93 | 58.1 |

**LULC\*- Land Use Land Cover**

**Table 8:** Pearson Correlation analysis.

| Factors |  | Population growth | Forest cover change | Year of change |
| --- | --- | --- | --- | --- |
| Population growth | Pearson Correlation | 1 | -0.998* | 0.999* |
|  | Sig. value |  | 0.043 | 0.020 |

### Conclusion and Recommendations Conclusion

Forests play a critical role in maintaining an environment that supports sustainable development and contributes to the national economy. Beyond their positive impacts on climate conditions, forests are essential in controlling soil erosion, combating land degradation and desertification, and preserving biodiversity. This study analyzes the deforestation of the Yayo Coffee Forest Biosphere Reserve in relation to the link between population growth and changes in forest cover.

The data analyzed for this research were sourced from two key areas: survey assessments and satellite imagery. Both types of analysis confirmed that population increase is a primary driver of forest degradation. Other significant factors include the lack of local community participation, illiteracy, poverty, certain investments (particularly coal mining at the Yayo Fertilizer Factory), and infrastructure development such as road construction. Forest product collection—especially land clearing for coffee expansion—and agricultural land and settlement expansion were identified as the major ways in which population growth impacts the forest.

The Yayo Coffee Forest Biosphere Reserve has been recognized as an in-situ conservation area for wild Coffea arabica populations, based on its presence of wild coffee, its accessibility for research, and its rich faunal and floral biodiversity. Additionally, the reserve provides significant environmental services globally. Deforestation in this reserve would result in the loss of a wide array of faunal and floral diversity, as well as the disruption of climate regulation functions provided by the forest.

It is crucial that government bodies at all levels give due attention to mitigating—and, if possible, preventing factors contributing to deforestation, such as the expansion of agricultural land, land clearing for coffee production, and investments that do not prioritize local community development. Sustainable management practices that balance conservation and development are essential for preserving the forest’s ecological integrity and ensuring its long-term benefits.

### Recommendation

Here are the recommendations for improving the conservation and management of the Yayo Coffee Forest Biosphere Reserve based on the study’s findings:

- Encourage Local Community Participation: Actively involve the local community in the development and implementation of management and conservation plans for the biosphere reserve. Ensuring that local people are part of the decision-making process can help improve compliance and effectiveness.
- Promote Coffee Cultivation on Private Land: Encourage farmers to cultivate coffee on their own land or gardens rather than exploiting wild coffee diversity in the forest. This can help reduce pressure on the forest while allowing local communities to continue benefiting from coffee production.
- Empower Community-Based Customary Institutions: Strengthen and empower community-based customary institutions, rather than marginalizing them, to manage and conserve the biosphere. A real bottom-up approach to conservation that integrates indigenous knowledge and local governance systems will help reduce conflicts and promote sustainable forest management.
- Conduct Comprehensive Impact Assessments of the Extractive Industry**: Undertake** detailed research on the impacts of extractive industries, such as coal mining, on biodiversity, land, water resources, and community health. This should include assessments on heavy metal contamination and strategies for mitigating negative effects, particularly as the Yayo Fertilizer Factory’s coal mining operations expand.
- Address Conflicts through Governance Improvements: Review and address the rules governing the coffee forest protected area, particularly those that have been identified as sources of conflict. Work towards fair, inclusive policies that balance conservation needs with the livelihoods of local communities.
- Develop Alternative Livelihood Strategies: Identify alternative sources of income or benefits from the coffee forest that could mitigate the negative impacts of current land use practices. Providing sustainable livelihoods could reduce reliance on forest exploitation and address underlying issues that lead to conflicts.
- Future Research on Wild Coffee and Biodiversity: Future studies should explore the role of wild coffee systems in supporting forest biodiversity and demonstrate how forest coffee-based livelihoods and biodiversity conservation can complement each other, rather than be viewed as conflicting goals.
- Promote Free, Prior, and Informed Consent: Ensure that the local community gives free, prior, and informed consent before any national or international resource management rules are enforced. Integrating customary management systems into broader conservation strategies will help protect both local traditions and forest resources.
- Establish Clear Guidelines for Resource Management: Develop and implement clear guidelines that govern the behavior of both resource users and managers. These guidelines should promote effective conservation and sustainable use of the coffee forest, ensuring that all stakeholders understand their roles and responsibilities.
- Government Intervention on Investment and Land Displacement: The government should create boundaries for investments that could displace local communities from their land. Policies should be reconsidered to ensure that local communities benefit from development and are not unfairly displaced. Incentives for community development should be built into investment projects, ensuring that both environmental and social concerns are addressed.

These recommendations aim to foster a more sustainable, inclusive, and equitable approach to forest conservation in the Yayo Biosphere Reserve, balancing environmental protection with the well-being of local communities.

## Acronym/Abbreviations

CBOs: Community Based Organization
DAs: Developmental Agents
EARO: Ethiopian Agricultural Research Organization
FAO: Food and Agricultural Organization
IUCN: International Union for Conservation of Nature and Natural Resources
M.a.s.l: Meter above sea level
MAB: Program on Man and the Biosphere
NGOs: Non-Governmental Organization
SPSS: Statistical Package for Social Sciences
STCP: Sustainable Tree Crops Program
UNESCO: United Nations Educational, Scientific and Cultural Organization
ZEF: Zentrum fur Entwicklungsforschung (Center for Development Research)

## Data Availability statement

The original contributions of the study are provided in the article/Supplementary Material; any additional inquiries can be directed to the corresponding author.

## Author Contributions

Conceptualization, F.M. and A.A. and T.J.; methodology, F.M. and T.J.; software, T.J.; formal analysis, F.M.; writing—original draft preparation, T.J. and F.M; writing—review and editing, A.A. All authors have read and agreed to the published version of the manuscript.

## Institutional Review Board Statement

Not applicable.

## Data Availability Statement

The raw data supporting the conclusions of this article will be made available by the authors on request.

